# Artificial selection on male size depletes genetic variance but not covariance of life history traits in the yellow dung fly

**DOI:** 10.1101/664326

**Authors:** WU Blanckenhorn, V Llaurens, C Reim, Y Teuschl, E Postma

## Abstract

The evolutionary potential of organisms depends on the presence of sufficient genetic variation for traits subject to selection, as well as on the genetic covariances among them. While genetic variation ultimately derives from mutation, theory predicts the depletion of genetic (co)variation under consistent directional or stabilizing selection in natural populations. We estimated and compared additive genetic (co)variances for several standard life history traits, including some for which this has never been assessed, before and after 24 generations of artificial selection on male size in the yellow dung fly *Scathophaga stercoraria* (Diptera: Scathophagidae) using a series of standard half-sib breeding experiments. As predicted, genetic variances (V_A_), heritabilities (*h*^*2*^) and evolvabilities (I_A_) of body size, development time, first clutch size, and female age at first clutch were lower after selection. As independent selection lines were crossed prior to testing, we can rule out that this reduction is due to genetic drift. In contrast to the variances, and against expectation, the additive genetic correlations between the sexes for development time and body size remained strong and positive (*r*_A =_ 0.8–0.9), while the genetic correlation between these traits within the sexes tended to strengthen (but not significantly so). Our study documents that the effect of selection on genetic variance is predictable, whereas that on genetic correlations is not.

## INTRODUCTION

The evolutionary potential of organisms depends on the presence of sufficient genetic (co)variation for any trait subject to selection. However, genetic variation is not constant over time (Crow, 2008). While increases in genetic variation ultimately derive from mutation, evolutionary theory predicts the depletion of genetic variation in natural populations under consistent directional or stabilizing selection (Falconer, 1989; Roff, 1997; Lynch and Walsh, 1998; Bürger and Gimmelfarb, 1999; Zhang and Hill, 2002; Crow, 2008), and there are a number of classic examples of such effects in the literature (summarized in Chapter 12 in Falconer, 1989; Chapter 4 in Roff, 1997). Nevertheless, genetic variation is abundant for many life history traits related to fitness. A number of evolutionary mechanisms, such as mutation-selection balance, antagonistic pleiotropy, frequency-dependent selection, genotype-environment interactions, heterogeneous environments, variable or sexually antagonistic selection have been uncovered, all of which may contribute to the resolution of this apparent inconsistency (Chapter 9 in Roff, 1997; Lynch and Walsh 1998; e.g. Turelli and Barton, 2004; Coltman et al., 2005; Förster et al., 2007). Nevertheless, a general answer to what maintains genetic variation in the face of selection remains elusive, and available data are inconsistent, so more research on the issue is needed.

While the predicted depletion of genetic variation in response to stabilizing or directional selection is straightforward at least in theory, the effect of selection on genetic covariation (i.e. on genetic correlations) is not so clear (Chapter 5 in Roff, 1997; e.g. Czesak et al., 2006). There are essentially two types of genetic correlations: genetic correlations among various traits within individuals, and genetic correlations of the same trait across different environments or the sexes. The latter are of particular interest in research on sexual dimorphism (Lande, 1980; Meagher, 1992; Reeve and Fairbairn, 1996; Ashman and Majetic, 2006; reviewed by Poissant et al., 2010). Sexual size dimorphism (SSD) evolves by sexually antagonistic natural or sexual selection when the optimal body size of females and males differs (the differential equilibrium hypothesis: Fairbairn, 1997; Blanckenhorn, 2000, 2007). However, because most genes are localized on the autosomes and are consequently shared by the sexes, many traits will be strongly (positively) correlated between the sexes. The evolution of SSD therefore will be constrained to some extent by high inter-sexual genetic correlations (Lande, 1980), be they generated by the same genes (so-called intra-locus sexual conflict) or different genes (inter-locus sexual conflict; Rice and Chippindale, 2001; Day and Bondurianski, 2004; see also Fairbairn and Roff, 2006). In general, the evolution of SSD by sexually antagonistic selection requires and therefore predicts the breakdown of such intersexual genetic correlations (Lande, 1980; Reeve and Fairbairn, 1996; Poissant et al., 2010). Genetic correlations among traits within the sexes are also likely to change under strong directional selection, although their evolution depends on the circumstances and is not easily predictable (Roff, 1997; Czesak et al., 2006).

In this study we estimate and compare genetic variation and covariation for several life history traits before and after artificial selection in the yellow dung fly *Scathophaga stercoraria* (Diptera: Scathophagidae). In the context of body size and SSD evolution, we artificially selected upwards and downwards on male body size for 24 generations. Selection in- or decreased body size by ca. 10% and no plateau was reached (Teuschl et al., 2007). Here we investigate sex-specific body size, development time, age at first reproduction, first clutch size (females only) and lifespan, although only the first four traits were similarly assessed before and after selection. Genetic estimates for some of these traits in this species have been reported before (Blanckenhorn, 2002; Blanckenhorn and Heyland, 2004), but others not. We expected that directional artificial selection would deplete genetic variation to some extent. As body size selection was performed only on males, we further expected genetic correlations between the sexes to decrease in magnitude too, because males and females presumably have different optima in this dimorphic species (Blanckenhorn, 2000, 2007).

## MATERIALS AND METHODS

### Genetic variation before selection

On four different days in April and May 2000, a total of 120 copulating pairs of *Scathophaga stercoraria* were randomly collected at a pasture in Fehraltorf, Switzerland. Females were brought to the laboratory and provided with a smear of dung into which they could lay eggs. These F1 eggs were used to start two replicate sets of flies, each stemming from at least 50 females, of which the offspring were raised under the standard laboratory conditions given below. For each of these sets, 50 males and 50 females from as many F1 families as possible were then randomly assigned to one of two small (S), large (L), or control (C) body size selection lines. The selection procedure, described in detail in Teuschl et al. (2007), started with the F2 offspring of these six selection lines. Body size truncation selection (selection differential ∼1 standard deviation per generation) was performed on males only.

Two subsets of our half-sib/full-sib/two container design before selection, as described below, were performed using F2 laboratory offspring of the two replicate control (C) lines. The third subset of our initial design was conducted with the F1 offspring of a further set of field flies collected in Fehraltorf later in the year 2000.

### Genetic variation after selection

Selection was first performed for 21 generations (approx. 2.5. years). At generation 21, flies from the two replicates of each selection regime (S, L, C) were crossed to control for potential inbreeding effects and to restore heterozygosity that might have become depleted by genetic drift during the selection process (Teuschl et al., 2007). Selection was further continued with these crossed S, L and C lines until generation 24. Thereafter the flies were propagated without selection for two more generations, at which point (in 2003) assessment of genetic (co)variation after selection took place using a half-sib/full-sib design analogous to that before selection.

### General rearing design

For our estimation of genetic variance and covariance we performed classic half-sib/full-sib/two container designs as outlined in Tables 2.4 & 2.5 by Roff (1997). Before and after selection, we mated three subsets of originally 10 and 12 males (sires) each to 3 females (dams), to produce a total of 76 (27 sires) and 97 (35 sires) full-sib offspring families before and after selection respectively (some families dropped out *a posteriori*). To separate common environment (container) effects from maternal effects, each full-sib family was split into two rearing containers (Roff, 1997). Before selection, the three subsets stemmed from the two control plus the later field-collected flies as described above (generation 2). After selection, the three subsets stemmed from the crossed S(mall), L(arge) and C(ontrol) lines at generation 26. In all cases, pairs were formed randomly within the subsets (avoiding full-sib crosses). Final family sizes (i.e. offspring numbers), from which all data were derived, varied between 2 and 15 per sex per dam and between 8 and 32 per sire.

For rearing, 10-25 eggs from each clutch were transferred into 100 ml plastic containers with an overabundant amount of ca. 80 g cow dung (= larval food). The flies were reared at constant 20 °C, 60 % humidity and 12 h photoperiod until offspring emerged after 19-22 days. Emerging adult flies were kept singly in 100 ml bottles at the same environmental conditions with water, sugar and *Drosophila melanogaster ad libitum* as prey until their death. Yellow dung flies are nutritionally anautogenous income breeders, as adult flies of both sexes require prey to produce gametes and to become sexually mature (Blanckenhorn and Henseler, 2005). Adult males were tested daily for sexual maturity (i.e. successful copulation), starting at age 3 days, with females aged 5 days or older.

### Traits assessed

For each individual offspring we assessed the following sex-specific traits (females only for all fecundity traits): hind tibia length as a practical surrogate of body size in this species; egg-to-adult development time; adult age at first copulation; first clutch size (i.e. egg number); female adult age at first clutch (first reproduction); adult lifespan in the holding bottle; lifetime egg number; eggs laid per day (= lifetime egg number / (adult lifespan – age at first clutch)). Genetic parameters for some of these traits, as well as some correlations among them, were reported previously for this species derived from other experiments (Blanckenhorn, 2002; Blanckenhorn and Heyland, 2004), but some traits (lifespan, first copulation) have never been assessed in a quantitative genetic context, so they are reported here for the data set before selection only. Only four of these traits (body size, development time, first reproduction, female first clutch size) were assessed in the same manner before and after selection, and therefore constitute the basis of our main comparison.

### Quantitative genetic analyses

Despite the classic half-sib/full-sib breeding design employed, quantitative genetic parameters were estimated by fitting an animal model (e.g. Kruuk, 2004; Thompson, 2008; Hill and Kirkpatrick, 2010; Wilson et al., 2010). This has the advantage of providing a direct estimate of the causal variance components, rather than the observational sire and dam variances. All mixed model analyses were performed in ASReml 4.1 (Gilmour et al., 2015).

In addition to the additive genetic animal effect, providing an estimate of the additive genetic variance (V_A_), we estimated the variance explained by maternal ID (V_M_), which captures the resemblance among full sibs over and above the resemblance expected based on additive genetic effects. We additionally fitted container ID to account for any additional resemblance among individuals that grew up in the same container (V_C_; ‘before selection’ only). The only fixed effects included were a trait-specific intercept and, when replicates were combined (see below), replicate.

Initially, homologous traits measured in both males and females were treated as different traits, as were the same traits measured before and after selection. Variance components were estimated simultaneously for all 18 ‘traits’ (8 female and 4 male traits before selection, 4 female and 2 male traits after selection) by fitting multivariate animal models with diagonal covariance structures (i.e. covariances were not estimated).

Models were fitted to the raw trait values, as well as to variance-standardised and mean-standardised trait values. The latter was done by dividing trait values by their standard deviation and mean, respectively, allowing us to directly estimate narrow-sense heritabilities (*h*^2^) and evolvabilities (I_A_) for all traits. Note that I_A_ relates to the coefficient of additive genetic variance, CV_A_, by I_A_ = (CV_A_/100)^2^ (Houle, 1992). Although CV_A_ may be a more familiar measure of variation, I_A_ is preferable because its numerical value has a more direct interpretation (Hansen and Houle, 2008). These transformations permit comparisons among traits with different means and variances, and also account for any differences in the mean or variance before and after selection.

As the breeding experiment was replicated thrice, in a first step we estimated V_A_, *h*^2^ and I_A_ for each replicate separately. However, as sample sizes (i.e. number of sires) per replicate were small, replicate-specific variance components were estimated with great uncertainty. We compared the log-likelihood of these models, estimating V_A_, *h*^2^ or I_A_ for each replicate, to models in which estimates were constrained to be the same across replicates. Note that in the latter models we left the remaining random effects unconstrained, as they were not of direct interest here. Twice the difference in log-likelihood of the two models was assumed to be approximately χ^2^-distributed with 36 degrees of freedom. In none of these comparisons the more complex model, estimating replicate-specific parameters, was significantly better than the constrained model (V_A_ : χ^2^ = 19.4, P = 0.99; *h*^2^: χ^2^ = 16.6, P = 1.00; I_A_ : χ^2^ = 20.9, P = 0.98). Hence we concluded there is no evidence that estimates of V_A_, *h*^2^ or I_A_ differed among replicates.

For all subsequent analyses replicates were combined to obtain an overall estimate for each trait, reducing the number of estimable variance components from 54 to 18. V_A_, *h*^2^ and I_A_ for all 8 female and 4 male traits measured before selection, and the 4 female and 2 male traits measured also after selection, were estimated using a model similar to that outlined above, but now homologous traits measured in different replicates were treated as the same trait. In addition to an additive genetic, a maternal and a container effect (before selection only), and a trait specific intercept, we fitted replicate as an additional fixed effect to account for any differences in the mean trait value among replicates.

Although in all analyses homologous traits measured in males and females were treated as different traits, we did test for sex-differences in V_A_, *h*^*2*^ and I_A_ for the six traits measured in both sexes (before: body size, development time, age at first copulation and lifespan; after: body size and development time) by comparing a model in which V_A_, *h*^*2*^ or I_A_ were constrained to be the same in both sexes using a likelihood-ratio test with six degrees of freedom.

Having obtained estimates of V_A_, *h*^*2*^ and I_A_ for males and females before and after selection, we tested whether any of these quantitative genetic parameters had changed during selection. This was done by comparing the log-likelihood of this (unconstrained) model to a model in which variances for male and female body size and development time, as well as for female age at first clutch and clutch size, were constrained to be the same before and after selection. Again, both models were compared using a likelihood-ratio test with six degrees of freedom.

Finally, we tested whether either intra- or inter-sexual additive genetic correlations differed before and after selection. As it was computationally impossible to simultaneously estimate all pairwise correlations, we restricted ourselves to a series of bivariate analyses of body size and development time, which were the only traits that had non-zero estimates of V_A_ (see Results) and were both measured in males and females before *and* after selection. To this end, we fitted a multivariate model to the two traits of interest (either male or female development time and size, or male and female development time or size) before and after selection (i.e. four traits in total). Correlations among traits measured before and after selection were set to zero. Similarly, the residual correlation between the sexes does not exist and was set to zero. Again, we used likelihood ratio tests with one degree of freedom to test whether this model provided a significantly better fit to the data than a model in which the additive genetic correlation before and after selection was constrained to be the same. In all these models the variances were always left unconstrained.

## RESULTS

### Genetic variance before and after selection

Estimates of V_A_, *h*^2^ and I_A_ with their approximate standard errors are provided in Table 1, and the other variance components (maternal, container and residual) are provided in Supplementary Table S1. Before selection, both male and female development time and body size showed substantial levels of additive genetic variance, as did age at first clutch and clutch size in females, and age at first copulation in males. The heritability of the life history traits was intermediate (*h*^*2*^ > 0.4), while that of male age at first copulation, a behavioural trait, was lower (*h*^*2*^ ≈ 0.25). These estimates are roughly in line with previous estimates for this species (Blanckenhorn, 2002; Blanckenhorn and Heyland, 2004) and with values for these traits for other species (Mousseau and Roff, 1987; Roff, 1997; Stirling et al., 2002; Table 1). In contrast, there was little to no additive genetic variance in female age at first copulation, male and female lifespan, and the compound measures of female reproductive output (total eggs and eggs per day). There was no evidence for sex-differences in V_A_ (χ^2^ = 3.58, P = 0.73), *h*^2^ (χ^2^ = 2.30, P = 0.89) or I_A_ (χ^2^ = 3.24, P = 0.78).

**Table 1.** Trait means, phenotypic variance (V_P_), additive genetic variance (V_A_), narrow-sense heritability (*h*^2^) and evolvability (I_A_) with their approximate standard errors (SE) for females (F) and males (M) before (B) and after (A) selection. Significant estimates with a *z*-ratio (estimate divided by standard error) greater than or equal to 1.96 are underlined.

| Trait | Sex | Mean | $V_p$ | $V_A \pm SE$ | $h^2 \pm SE$ | $I_A \pm SE (\times 10^3)$ |
| --- | --- | --- | --- | --- | --- | --- |
| **Before selection** |  |  |  |  |  |  |
| *development time (days)* | F | 23.6 | 3.07 | <u>$2.39 \pm 0.20$</u> | <u>$0.777 \pm 0.064$</u> | <u>$4.30 \pm 0.35$</u> |
| *size (mm)* | F | 3.01 | 0.0194 | $0.0113 \pm 0.0070$ | $0.583 \pm 0.362$ | $1.25 \pm 0.77$ |
| *age at 1<sup>st</sup> clutch (days)* | F | 10.8 | 5.04 | $2.19 \pm 1.41$ | $0.434 \pm 0.280$ | $18.9 \pm 12.2$ |
| *clutch size (eggs)* | F | 65.2 | 125 | <u>$57.5 \pm 17.7$</u> | <u>$0.459 \pm 0.141$</u> | <u>$13.5 \pm 4.2$</u> |
| *age at 1<sup>st</sup> copulation (days)* | F | 6.22 | 0.257 | $0.0043 \pm 0.0270$ | $0.017 \pm 0.105$ | $0.11 \pm 0.70$ |
| *total eggs laid* | F | 229 | 16911 | 0 | 0 | 0 |
| *egg production per day* | F | 7.03 | 12.1 | 0 | 0 | 0 |
| *lifespan (days)* | F | 48.2 | 490 | 0 | 0 | 0 |
| *development time (days)* | M | 25.7 | 4.08 | <u>$2.48 \pm 0.19$</u> | <u>$0.607 \pm 0.047$</u> | <u>$3.76 \pm 0.29$</u> |
| *size (mm)* | M | 3.67 | 0.0306 | <u>$0.0178 \pm 0.0014$</u> | <u>$0.581 \pm 0.045$</u> | <u>$1.32 \pm 0.10$</u> |
| *age at 1<sup>st</sup> copulation (days)* | M | 5.79 | 4.03 | $0.954 \pm 0.769$ | $0.237 \pm 0.191$ | $28.5 \pm 23.0$ |
| *lifespan (days)* | M | 70.7 | 1103 | 0 | 0 | 0 |
| **After selection** |  |  |  |  |  |  |
| *development time (days)* | F | 20.6 | 1.37 | $0.765 \pm 0.539$ | $0.560 \pm 0.394$ | $1.81 \pm 1.28$ |
| *size (mm)* | F | 2.88 | 0.0368 | $0.0029 \pm 0.0074$ | $0.078 \pm 0.201$ | $0.35 \pm 0.89$ |
| *age at 1<sup>st</sup> clutch (days)* | F | 14.5 | 6.43 | $0.934 \pm 1.832$ | $0.145 \pm 0.285$ | $4.42 \pm 8.66$ |
| *clutch size (eggs)* | F | 68.5 | 133 | $27.5 \pm 19.5$ | $0.207 \pm 0.147$ | $5.85 \pm 4.15$ |
| *development time (days)* | M | 22.1 | 1.31 | $0.160 \pm 0.333$ | $0.122 \pm 0.254$ | $0.33 \pm 0.68$ |
| *size (mm)* | M | 3.45 | 0.0465 | 0 | 0 | 0 |

For all traits that were measured before and after selection, V_A_, *h*^2^ and I_A_ were lower after selection (sign-test: P = 0.031), with narrow-sense heritabilities declining by 28% to 100% (on average 69%; Table 1). Nevertheless, as variances of individual traits before and after selection were estimated with rather high uncertainty (featuring large standard errors), explicit tests for differences in V_A_ before and after selection did not quite reach statistical significance (χ^2^ = 11.6, d.f. = 6, P = 0.072). The same was true when comparing heritabilities (χ^2^ = 10.34, P = 0.11) and evolvabilities (χ^2^ = 11.06, P = 0.087).

### Genetic correlations

Although the additive genetic correlation between female development time and size was weakly negative before (*r*_A_ = −0.14 ± 0.36) and positive after selection (*r*_A_ = 0.32 ± 0.89), the correlations were not significantly different from each other (χ^2^ = 0.15, d.f. = 1, P = 0.70). For males, the additive genetic correlation between development time and size before selection was also weakly negative (*r*_A_ = −0.043 ± 0.055), whereas after selection this correlation approached 1. However, the latter was poorly estimated due to lack of additive genetic variance in male size (see Table 1), so these correlations again did not differ significantly (χ^2^ = 0.27, d.f. = 1, *P* = 0.61).

The inter-sexual correlations between male and female development time were strong and positive both before (*r*_A_= 0.88 ± 0.08) and after selection (*r*_A_ = 0.91 ± 0.22), and not significantly different (χ^2^ = 0.0, d.f. = 1, P = 1.0). The same was true for the correlation between male and female size before selection (*r*_A_= 0.81 ± 0.10), but again due to the lack of additive genetic variance for male size after selection, the correlation between male and female size after selection was poorly estimated and bound at 1. In line with this, the two correlations were not significantly different (χ^2^ = 0.36, d.f. = 1, *P* = 0.55).

## DISCUSSION

We here compared additive genetic variances and correlations before and after 24 generations of artificial selection on male body size over a period of ca. 2.5 years in the yellow dung fly *Scathophaga stercoraria* (Teuschl et al., 2007). Thereby we could experimentally test the theoretical prediction that directional selection depletes genetic (co-)variation (Falconer, 1989; Roff, 1997; Lynch and Walsh, 1998; Bürger and Gimmelfarb, 1999; Zhang and Hill, 2002; Crow, 2008). Investigating four life history traits (body size, development time, first clutch size, female age at first clutch), we found indeed that, on average, additive genetic variation V_A_, heritabilities *h*^*2*^ and evolvabilities I_A_ were lower after selection than before (Table 1). This effect was mediated by a decrease in the narrow-sense genetic variance, as the environmental (i.e. residual) variance remained roughly constant or even increased (Table S1). Genetic variation was generally similar for both sexes (cf. Förster et al., 2007). Although we did not monitor neutral variation in this study (e.g. by way of microsatellites), we do not believe that this reduction of genetic variation was caused by random loss of alleles due to genetic drift in our (necessarily small) laboratory populations, as we conducted our post-selection assessment after crossing replicate selection lines to offset potential inbreeding and to restore any such loss of heterozygosity (Teuschl et al., 2007). Building on the classical studies showing or suspecting such an effect (see Falconer, 1989; Roff, 1997; Lynch and Walsh, 1998), here we thus provide an experimental test confirming theoretical predictions of depleting genetic variation in response to directional (or stabilising) selection,.

Theory predicts a breakdown of typically strongly positive inter-sexual genetic correlations in response to selection favouring sexual dimorphism (Lande, 1980; Meagher, 1992; Reeve and Fairbairn, 1996, 2001). In contrast to the depletion of genetic variation, we however found no change in the magnitude of the additive genetic correlation between the sexes in development time and body size: these correlations roughly remained at their initially high level of *r*_A =_ 0.8 – 0.9. Yellow dung flies are sexually dimorphic (males larger), and the inter-sexual genetic correlation for body size and associated traits (such as development time) is indeed less than one (Blanckenhorn, 2002, 2007; this study). However, artificial selection did not further reduce this correlation beyond the level present before selection. We only artificially selected on male size, and our selection procedure resulted in a strong parallel response in female size (Teuschl et al., 2007). We suspect that the time frame of our selection experiment was simply too short to obtain a change, as Lande’s (1980) models suggest that breaking down genetic correlations between the sexes takes considerable time (see also Reeve and Fairbairn, 1996, 2001).

For the genetic correlations between size and development time within the sexes we however found the opposite: although this effect was not statistically significant, these correlations tended to become more positive. In yellow dung flies this genetic correlation is typically positive in various larval environments but weak on average (Blanckenhorn, 1998; Teuschl et al., 2007), contrary to the situation in *Drosophila melanogaster*, where it is strong and close to 1 (e.g. Nunney, 1996). The trend found here is thus opposite to the general prediction of a breakdown of genetic covariation in response to selection. Due to lack of power our conclusions in this regard here must remain limited. Nevertheless, it is congruent with mounting evidence that the evolution of genetic correlations in the face of selection and environmental change is actually not easily predictable (Roff, 1996, 1997; Czesak et al., 2006).

In addition to presenting an explicit test of the effect of directional selection on additive genetic variances and correlations, our study further provides genetic estimates for some life history and behavioural traits that have never before been documented for the yellow dung fly. These were assessed before selection, thus reflecting naturally present genetic variation (Table 1). Most of these traits (adult lifespan in the laboratory, lifetime fecundity measures, age at first copulation) had low, non-significant heritabilities. This may relate precisely to the effect documented here, as persistent selection especially and most strongly is expected to deplete the genetic variation of traits closely related to fitness (Falconer, 1989; Roff, 1997; Lynch and Walsh, 1998; Bürger and Gimmelfarb, 1999). However, while the lifespan of a single fly in a bottle should reflect its intrinsic longevity to some extent, natural selection is largely suspended in laboratory settings, so that random and unnatural mortality events may render such estimates of survivorship and lifetime fecundity doubtful if not meaningless.

In conclusion, we present an empirical test in the yellow dung fly of the theoretical prediction that genetic (co-)variation should become depleted in response to, in this case, artificial directional selection on body size (Teuschl et al. 2007). Across a number of life history and behavioural traits, we indeed found that genetic variation after selection was reduced. In contrast, initially strongly positive genetic correlations between the sexes did not become reduced, while the genetic correlation between development time and body size within the sexes actually became more positive on average, but not significantly so. We believe our results are not simply due to random loss of heterozygosity in small laboratory populations, as independent selection lines were crossed at the end prior to testing. Our data set adds new evidence to the traditional classic studies of this phenomenon (Falconer, 1989; Roff, 1997; Lynch and Walsh, 1998) and shows that the effect of selection on genetic variance appears predictable, while that on genetic correlations is not (Roff, 1997; Czesak et al., 2006).

## ACKNOWLEDGEMENTS

We thank the Swiss National Science Foundation for funding this project. V. Llaurens participated in this study during an internship at the University of Zurich set up by the INAPG, France.

---

**Table S1.**
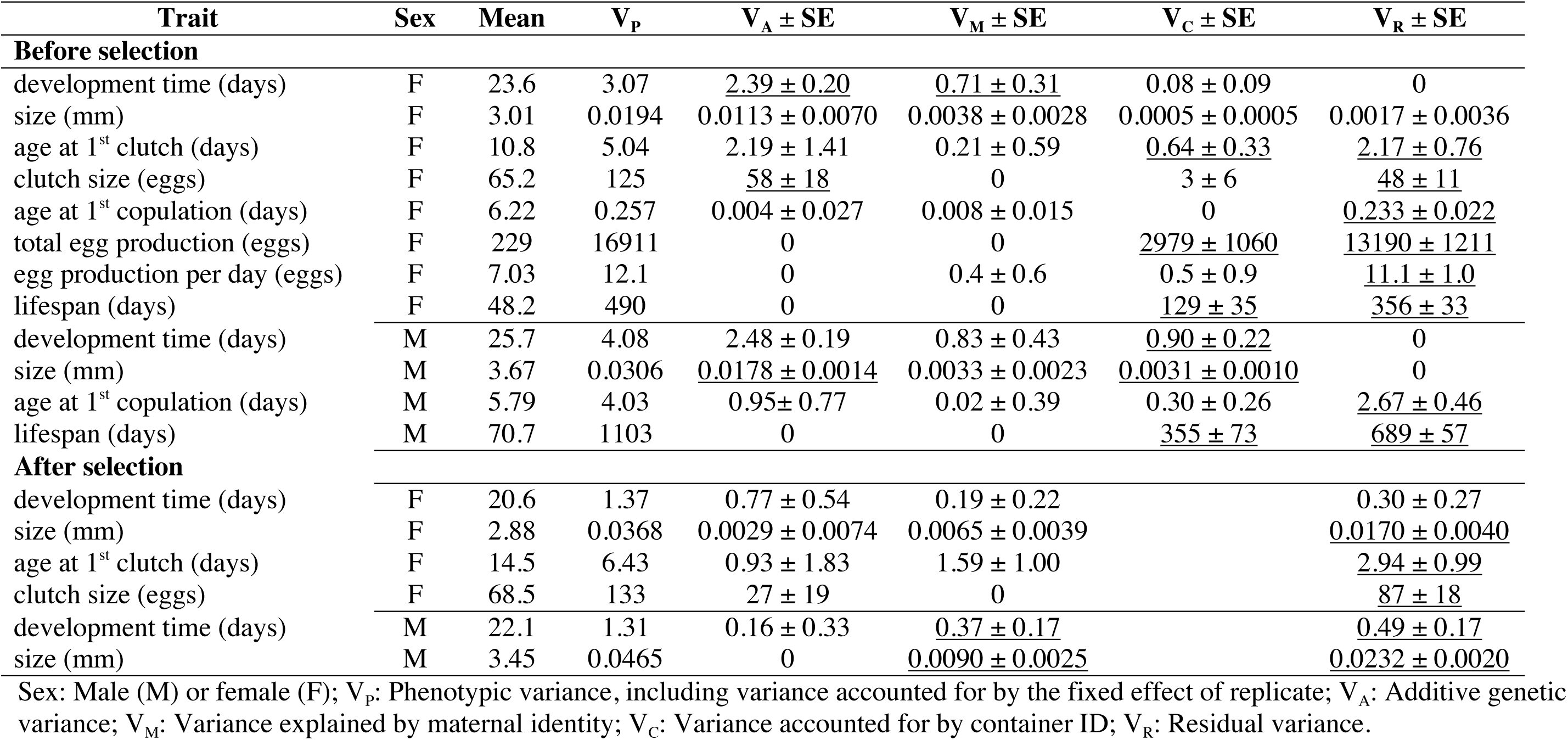
Variance components for all random effects ± their approximate standard error. Only estimates for unstandardised trait values are provided, as mean- and variance-standardised estimates can easily be obtained by dividing by the squared trait mean and V_P_, respectively. Estimates with a *z*-ratio greater than or equal to 1.96 are underlined. Note that V_P_ and the sum of V_A_, V_M_, V_C_ and V_R_ are not necessarily the same; this is because the former includes variance among replicates, which was accounted for by including replicate as a fixed effect in the animal models, and because variances were constrained to be positive.

## 5 anonymous peer review attached

### Referee #1

The paper by Blanckenhorn et al examines changes in genetic variance and correlations between traits for the yellow dung fly after 24 generations of selection for body size in males.

The experimental study and subsequent data collected are an excellent attempt at addressing an ongoing and interesting question about how genetic variances and correlations change under selection. Unfortunately, I don’t think the sample sizes the authors have been able to collect are sufficient to answer the questions posed. Further, I am a little confused by some of the analytical approaches used to circumvent this and I’m not sure I follow some of the reasoning and expectations. I go into some more detail below. It would have been nice if the authors had added page numbers, as it is it is a little hard to reference specific areas in the manuscript.

-At the end of page 4 the authors’ state that they expect the correlation to decrease between the sexes, but this will depend on the amount of sex-specific standing genetic variation and the strength of antagonistic selection. There is no measure of the strength of selection on females, so there is no way to know in each selection treatment (small vs large) where the female optimum lies. The lack of change in the genetic correlation between males and females in this experiment could result from either a lack of sex-specific variation or because there is no (or very weak) selection on female size to counter that of the artificial selection on males (sex-specific selection).
-The comparison of interest in selection studies is between the control (any differences between the control and the starting population would be due to the experimental set up, sampling and drift) and the selection treatments. It is a shame that the authors mixed the replicates up, rather than using the controls to account for these differences, thus adding much needed replication to some of these comparisons. The control lines can thus serve two purposes in selection experiments like this. First, they allow the removal of the effects of temporal fluctuations of means that can occur during selection experiments, (this is done by subtracting the average of the means of the control lines after selection from the means of the selection lines). Second, the control lines allow a comparison of the behavior of male and female trait (co)variance when directional selection is prevented from acting, but still having experienced the same experimental conditions. If genetic variance has decreased in the control population after selection, then you cannot rule out drift/sampling causing the decrease in variance between the original population and the selection populations.
-Carrying on from the point above, I do not follow the logic of the approach in the mixing of the selection treatments (including the control) for analysis and subsequent interpretation and then predicting that variance would be expected to decline. I understand that due to sample size, there was a need to try to increase the number of sires and for the pre-selection estimates this could be easily justified (even then the number of sires is still very much on the low side). However, there are three selection treatments, small, large and a control. Although it is expected that genetic variance would decrease in either of the selection treatments, combined you have effectively created a population under disruptive selection and therefore genetic variance would be expected to increase. The authors attempt to address this by including treatment as a fixed effect (essentially zeroing the mean), but this doesn’t account for heterogeneity in variances among treatments. The statistical test for this is not sufficient to say there is no difference, simply because the sample sizes of the sires are so small. Furthermore, as mentioned above, there should be detectable differences between the control and the selection treatments at the end of the experiment, and by including the control in the “post-selection” group for analysis you should find no difference as the variance in the control group will only be different from the starting population due to factors other than artificial selection.

### Referee #2

This study addresses fundamental questions in evolutionary biology regarding the effects of sex-specific directional selection and, specifically, its role in depleting genetic variance and breaking genetic correlations between the sexes, which allows independent evolution to sex-specific optima (sexual dimorphism). These a priori predictions stem from a robust theoretical framework that has been subjected to relatively little empirical work and thus makes this an important topic of investigation. Using bi-directional selection on male body size (two replicates per treatment) for 24 generations, this study tests these predictions while also providing novel estimates of genetic variance for various life-history traits in *Scathophaga stercoraria*, an important model system with male-biased sexual size dimorphism. However, I have several questions and concerns about the selection procedure, assays used to measure traits, and the authors’ ability to properly address the aims of this study.

1. The a priori expectation that directional selection on male size should break down genetic covariance between the sexes assumes that males and females experience different selection pressures that favor different optimal body sizes. That is, selection may favor intermediate female size while sexual selection drives larger male size; however, in the current study, selection was only implemented on male size. Therefore, by removing selection on females and essentially ignoring the costs associated with increasing or decreasing body size, it is not clear that one should expect a breakdown in genetic covariance. Instead, it seems likely that female size would simply track changes in male size. This seems like an important point that warrants attention, as it may mean that the experimental design is inappropriate for testing this specific prediction.
2. As noted in the manuscript, the authors’ power to detect changes in (co)variance before and after selection is limited by the fact that there were only two replicates per treatment, which were then crossed to restore heterozygocity, further reducing statistical power. Therefore, parts of the current study seem like qualitative assessments (i.e., is the change in (co)variance in the expected direction?) rather than quantitative.
3. When males and females were assayed for their age at first copulation, how old were their mates? From the text, it is my understanding that males were tested with females aged at least 5 days; however, previous work in this species has demonstrated that young females (< 8 d old) can be unattractive to males and/or resist mating attempts (Blanckenhorn et al, 2007, Behavioural Ecology), which could confound this measure.
4. The Materials and Methods section talks about three selection treatments (small, control, and large); however, there are no mentions of treatment specific effects of the selection regimen on (co)variance in the Results (including Tables 1 and S1) or Discussion. For example, both tables show trait values before and after selection but do not specify from which treatment (i.e., selection line) these values were obtained. I would be interested to see these data reported separately for each of the different selection lines.

### Referee #3

In this article the authors investigate the effect of artificial, directional selection on the maintenance of genetic variance and co-variance in life-history traits of the yellow dung fly *Scathophaga stercoraria*. They estimate genetic (co)variance for the traits in question in males and females both before and after 24 generations of artificial directional selection on male body size. The authors find that such selection significantly depleted additive genetic variance for the traits but actually strengthened genetic co-variance (albeit non-significantly). In general, the paper is well-written and presents both novel and interesting results.

#### Major concerns -

My main concern with the paper is the clarity or lack thereof of the methods section which I found confusing to read and think needs reworking. I appreciate that the setup of the selection lines is complicated and there is a lot to get across to the reader, but feel that this must be worked on in order to keep the reader’s attention and understanding of the otherwise well-written paper.

Materials and Methods: -

1. Please provide some brief detail on how body size truncation selection was performed, it is not enough to say that the selection procedure is explained elsewhere (line 107).
2. The authors mention “later field-collected flies” several times but as far as I can see do not explain why these flies were collected separately or the purpose of these extra flies.
3. Please explain why only four of the traits in question were assessed in the same manner before and after selection (lines 158-160).
4. 4. Please explain why the container effect was measured before selection only (lines 206-108).

#### Minor concerns: -

Introduction: -

1. Lines 81-83: Please give an example or explanation as to how circumstances might affect how genetic correlations respond to selection.
2. Line 86: Mention here that S. stercoraria are sexually dimorphic.
3. Lines 87-88: This sentence doesn’t seem to be connected with the surrounding sentences. Please clarify or remove.
4. Lines 90-91: Again please explain why only the first four traits were similarly assessed before and after selection.
5. Line 92: Please specify for which traits genetic estimation in your study is novel.
6. Line 95: Why do males and females “presumably” have different optima for body size?

Materials and methods: -

7. Lines 105-107: This sentence is very confusing, the authors need to make it explicitly clear that there were two replicates of three selection lines.
8. Line 109: There is a bullet point before 1 standard deviation, was this supposed to be a different symbol?
9. Lines 110-113: I was not satisfied that this was explained later in the methods. Again, what was the purpose of the later collected set of flies?
10. Line 143: Please explain what an “anautogenous income breeder” is.

### Referee #4

I applaud the intent of the experiment. However, I think that the issue of genetic drift during the course of selection needs to be taken into account.

The present analyses do not separate the effects of drift from those of selection. Given the design it would be nice to see the change in realized heritability of the selected trait: this could be using a cumulative approach. Selection will not only deplete genetic variance but will also cause linkage disequilibrium which could result in changed (co)variance components. The separate estimates are not statistically independent and so care should be taken in interpreting the results. In particular would caution against placing too much attention on individual estimates. I suggest an analysis that enters before and after as a dummy variable, which would allow a direct test of changes in additive genetic variance. This can be done with the anova approach or with the animal model (note that asreml is now freely available and instructions for between environment analyses described on the wamwiki site).

I don’t like family means as estimates of rg and would prefer to see direct tests of the covariances/genetic correlations. Standardizing the data before analysis would remove problems of scale.

In conclusion, I think that is a very nice data set to address the question of changes in additive (co)variances but a more refined analysis is required. Having said this my feeling is that the overall results will not change, though I suspect that it will not be possible to differentiate the effects of drift from selection.

### Referee #5

This manuscript analyzes data from a previous selection experiment in the yellow dung fly that selected to either increase or decrease body size. Using a half-sib design, the authors used an analysis of variance approach to calculate heritabilities and genetic correlations for life history traits before selection and then following 24 generations of selection. They found that heritabilities generally decreased, and on average within-sex correlations increased and between-sex correlations decreased. The authors discuss their findings within the context of the evolution of sexual size dimorphism and the loss of genetic variation following selection.

On one level, I found this manuscript to be interesting - the authors have measured a large number of traits and conducted selection for a considerable number of generations, and as a result the dataset has the potential to address some interesting questions about how selection modifies the G matrix. However, on another level, I am worried about a few aspects of the author’s experimental design and analysis, and I think the authors need to consider more of the current literature in the interpretation and extension of their findings.

If I am understanding the methods correctly, the pre-selection genetic parameters were estimated using a dataset that included the two control lines from the selection experiment as separate blocks, and then a third block that was generated from a separate collection of parental flies from the same location in a different year. Then the post-selection genetic parameters were estimated from a dataset that included the selection line to increase body size, the selection line to decrease body size, and the control line as separate blocks. Because these two datasets are not simply the same population before and after selection, the comparison between them loses validity. The post-selection parameters may be different from the pre-selection parameters simply because they are different subsamples of the same population, not because selection had any effect.

In addition, I find it strange that the authors would lump the control lines, selection lines to increase body size, and selection lines to decrease body size into one dataset that generated one set of parameter estimates. If the point is to determine how selection changed the parameters, including the control lines that were not under selection dilutes the effect of selection. Also, combining data from both selection lines means that the authors are unable to detect whether selection in one direction affected variation differently than selection in the other direction, or even whether selection had any effect on the traits being observed. If the authors have enough individuals and families in each line, I suggest analyzing the large, control, and small lines separately, and then using the control line after 26 generations as the standard of comparison, rather than the current pre-selection dataset.

The authors are estimating genetic covariation and correlations among multiple traits across the sexes, which is essentially a genetic variance-covariance matrix (the G matrix) with sex-specific submatrices (Gm and Gf) and a between-sex submatrix (B), as originally described by Lande (1980, Evolution, vol. 34, p. 292). Yet the authors do not use this terminology, nor do they seem to view their work in a multivariate context, which could be a more powerful framework and connect their work more effectively to what others have discovered about the topic in recent years. The ANOVA/ANCOVA approach to estimating parameters is certainly valid and productive, but increasingly other authors are using the animal model (Thompson 2008, Proc Biol Sci. Mar 22;275(1635):679-86), which would allow the authors of this manuscript to include all pedigree information simultaneously instead of generating separate full-sib and half-sib estimates, and it would estimate the between-sex correlations directly, rather than from the family means, which tend to underestimate the correlations.

There are also many statistical tests for comparing matrices, some better than others. Using one of these tests would allow the authors to test their question of whether the matrix changed as a result of selection using the entire matrix, rather than collapsing all variation and terms into an average across traits (as in Figs. 1 and 3) that retains less biological meaning. With a test like that of Calsbeek and Goodnight (2009, Evolution, 63(10), 2627-2635), the authors could compare the matrices within the context of the multivariate breeder’s equation, which would have the added benefit of making their findings more biologically relevant, and allow them to speculate on the future evolution of individual traits. The authors should also consider connecting their work more directly to the work of others on the same subject, such as Delph et al. (2011, Evolution, 65(10), 2872-2880), the meta-analysis by Barker et al. (2010. Evolution 64:2601-2613), the review by Cox and Calsbeek (2009, The American Naturalist Vol. 173, No. 2 pp. 176-187), etc. A better review of the literature would allow the authors to present a more sophisticated introduction and discussion that addresses the possible mechanisms and evolutionary consequences of a changing matrix, and allow them to go beyond just observing what happened to explaining how it might have happened and why it matters.

I also have a few more specific observations/recommendations:

1. The authors use the term (co)variation somewhat liberally, when they should be using variation and covariation as separate terms in a more specific fashion; for example, in the first sentence of the abstract the use of (co)variation implies that covariation is required for the evolution of a trait, which is not quite true-a trait with sufficient variation that does not covary with any other traits is still able to evolve. As the authors themselves point out, our understanding of how variation changes under selection is very different from how covariation changes, so in the second sentence of the abstract it’s inappropriate to lump the two together with (co)variation. The authors need to be more careful with their use of the term, and specify the difference between variation and covariation more clearly.
2. It’s not apparent from this manuscript what measurement of body size was under direct selection in the initial experiment, and whether that trait was included in this analysis. These facts are definitely of interest to me, and would help with interpretation of the results; how much variation was there in the trait under selection initially? What other traits are most likely to be correlated with it, perhaps through developmental constraints? Was the trait under selection sexually dimorphic to begin with, and if so, how much?
3. I found the description of the selection lines and pedigree structure to be somewhat unclear; I had trouble figuring out the difference between lines vs. subsets vs. blocks, for example, and it was confusing to have the half-sib/full-sib/container design presented a page before it was actually described. Perhaps a diagram would be of assistance here.
4. How was the significance testing for the correlations in Table 2 conducted? And why are some values in italics in Table 2?
5. The authors do not present all of the heritabilities and correlations they estimated in the post-selection analysis. I would like to see these presented in a table; generating estimates of genetic covariances is a lot of work, and as our understanding of the evolution of G changes, and potentially as other authors explore the evolutionary genetics of this species further, it could be valuable to have these estimates available in the literature.
6. In the second paragraph of the discussion, third sentence, the authors seem to have a misconception about between-sex genetic correlations. For the same trait measured separately in both sexes, the between-sex correlation can be one for a trait that is dimorphic, even strongly so. If the loci contributing to dimorphism are fixed at a particular allele, they will be contributing to the difference between the sexes but not contributing to the genetic variation in the trait. And the correlation only captures covariation, not absolute difference. A genetic correlation of 1 means that all of the genetic variation in the trait is shared across the sexes; it can’t tell us anything about how different the two sexes actually are in absolute terms.
7. Later in the third paragraph of the discussion, the authors speculate on why artificial selection reduced between-sex genetic correlations, but their logic of why selection only on one sex would cause the males and females to “drift further apart” is somewhat unclear, and it’s also unknown whether selection actually did change the absolute difference between males and females, or if correlated change in females kept the magnitude of dimorphism the same (or even if the trait under direct selection in the original experiment is the same as the proxy for body size measured in this experiment). I’d like to see their proposed mechanism for change explained in more detail.

Overall, I think the dataset generated for this manuscript has the possibility to be interesting and make a useful contribution to our understanding of how the G matrix changes under selection, but I believe that the authors need to generate estimates from the selection lines separately, update their analysis techniques to include matrix comparison methods and a multivariate approach, and incorporate more of the current literature in the introduction and discussion.

## References

Ashman TL, Majetic CJ (2006). Genetic constraints on floral evolution: a review and evaluation of patterns. Heredity 96: 343–352.

Blanckenhorn WU (1998). Adaptive phenotypic plasticity in growth, development and body size in the yellow dung fly. Evolution 52: 1394–1407.

Blanckenhorn WU (2000). The evolution of body size: What keeps organisms small? Quart Rev Biol 75: 385–407.

Blanckenhorn WU (2002). The consistency of heritability estimates in field and laboratory in the yellow dung fly. Genetica 114: 171–182.

Blanckenhorn WU (2007). Case studies of the differential equilibrium hypothesis of sexual size dimorphism in dung flies. In: Fairbairn DJ, Blanckenhorn WU, Szekely T, (eds). Sex, Size and Gender Roles. Evolutionary Studies of Sexual Size Dimorphism. Oxford University Press: Oxford UK, pp. 106–114.

Blanckenhorn WU, Henseler C (2005). Temperature-dependent ovariole and testis maturation in the yellow dung fly. Entomol Exp Appl 116: 159–165.

Blanckenhorn WU, Heyland A (2004). The quantitative genetics of two life history trade-offs in the yellow dung fly in abundant and limited food environments. Evol Ecol 18: 385–402.

Bürger R, Gimmelfarb A (1999). Genetic variation maintained in multilocus models of additive quantitative traits under stabilizing selection. Genetics 152: 807–820.

Coltman DW, O’Donoghue P, Hogg JT, Festa-Bianchet M (2005). Selection and genetic (co)variance in bighorn sheep. Evolution 59: 1372–1382.

Crow JF (2008). Maintaining evolvability. J Genet 87: 349–353.

Czesak ME, Fox CW, Wolf JB (2006). Experimental evolution of phenotypic plasticity: how predictive are cross-environment genetic correlations? Am Nat 168: 323–335.

Day T, Bonduriansky R (2004). Intralocus sexual conflict can drive the evolution oof genomic imprinting. Ann Rev Ecol Syst 40: 103–125.

Fairbairn DJ (1997). Allometry for sexual size dimorphism: Pattern and process in the coevolution of body size in males and females. Ann Rev Ecol Syst 28: 659–687.

Fairbairn DJ, Roff DA (2006). The quantitative genetics of sexual dimorphism: assessing the importance of sex-linkage. Heredity 97: 319–328.

Falconer DS (1989). Introduction to Quantitative Genetics, 3rd edn. Longman Scientific and Technical: Harlow.

Förster K, Coulson T, Sheldon BC, Pemberton JM, Clutton-Brock TH, Kruuk LEB (2007). Sexually antagonistic genetic variation for fitness in red deer. Nature 447: 1107–1110.

Gilmour AR, Gogel BJ, Cullis BR, Welham SJ, Thompson R. 2015 ASReml user guide release 4.1 Structural Specification, VSN International Ltd., Hemel Hempstead, HP1 1ES, UK. www.vsni.co.uk.

Hansen TF, Houle D (2008). Measuring and comparing evolvability and constraint in multivariate characters. J Evol Biol 21: 1201–1219.

Hill WG, Kirkpatrick M (2010). What animal breeding has taught us about evolution. Annu Rev Ecol Evol Syst 41: 1–19.

Houle D (1992). Comparing evolvability and variability of quantitative traits. Genetics 130: 195–204.

Kruuk LEB (2004). Estimating genetic parameters in natural populations using the ‘animal model’. Phil Trans R Soc Lond B 359: 873–890.

Lande R (1980). Sexual dimorphism, sexual selection, and adaptation in polygenic characters. Evolution 34: 292–305.

Lynch M, Walsh B (1998). Genetics and Analysis of Quantitative Traits. Sinauer Associates: Sunderland MA.

Meagher TR (1992). The Quantitative Genetics of Sexual Dimorphism in *Silene latifolia* (Caryophyllaceae). I.Genetic Variation. Evolution 46: 445–457.

Mousseau TA, Roff DA (1987). Natural selection and the heritability of fitness components. Heredity 59: 181–198.

Nunney L (1996). The response to selection for fast larval development in *Drosophila melanogaster* and its effect on adult weight: an example of a fitness trade-off. Evolution 50: 1193–1204.

Puissant J, Wilson AJ, Coltman DW (2010). Sex-specific genetic variance and the evolution of sexual dimorphism: a systematic review of cross-genetic correlations. Evolution 64: 97–107.

Reeve JP, Fairbairn DJ (1996). Sexual size dimorphism as a correlated response to selection on body size: an empirical test of the quantitative genetic model. Evolution 50: 1927–1938.

Reeve JP, Fairbairn DJ (2001). Predicting the evolution of sexual size dimorphism. J Evol Biol 14: 244–254.

Rice WR, Chippindale AK (2001). Intersexual ontogenetic conflict. J Evol Biol 14: 685–693.

Roff DA (1996). The evolution of genetic correlations: an analysis of patterns. Evolution 50: 1392–1403.

Roff DA (1997). Evolutionary Quantitative Genetics. Chapman and Hall: New York.

Stirling DG, Reale D, Roff DA (2002). Selection, structure and the heritability of behaviour. J Evol Biol 15: 277–289.

Teuschl Y, Reim C, Blanckenhorn WU (2007). Correlated responses to artificial body size selection in growth, development, phenotypic plasticity and juvenile viability in yellow dung flies. J Evol Biol 20: 87–103.

Thompson R (2008). Estimation of quantitative genetic parameters. Proc R Soc Biol Lond B 275, 679–686.

Turelli M, Barton NH (2004). Polygenic variation maintained by balancing selection: Pleiotropy, sex-dependent allelic effects and G3E interactions. Genetics 166, 1053–1079.

Wilson AJ, Réale D, Clements MN, Morrissey MM, Postma E, Walling CA, Kruuk LEB, Nussey DH (2010). An ecologist’s guide to the animal model. J Anim Ecol 79: 13–26.

Zhang XS, Hill WG (2002). Joint effects of pleiotropic selection and stabilizing selection on the maintenance of quantitative genetic variation at mutation-selection balance. Genetics 162: 459–471.

